## Supplementary Information for "The last resort antibiotic daptomycin exhibits two independent antibacterial mechanisms of action"

Fig. S1 Effects of daptomycin on cell growth in *S. aureus* and *B. subtilis*

Fig. S2 Inhibition of L-form growth by daptomycin

Fig. S3 *B. subtilis* AK092 L-forms still synthesise lipid II, whilst AK0197 L-forms do not

Fig. S4 Overview images of Nile red-stained untreated *B. subtilis*

Fig. S5 Overview images of Nile red-stained daptomycin-lipid clusters

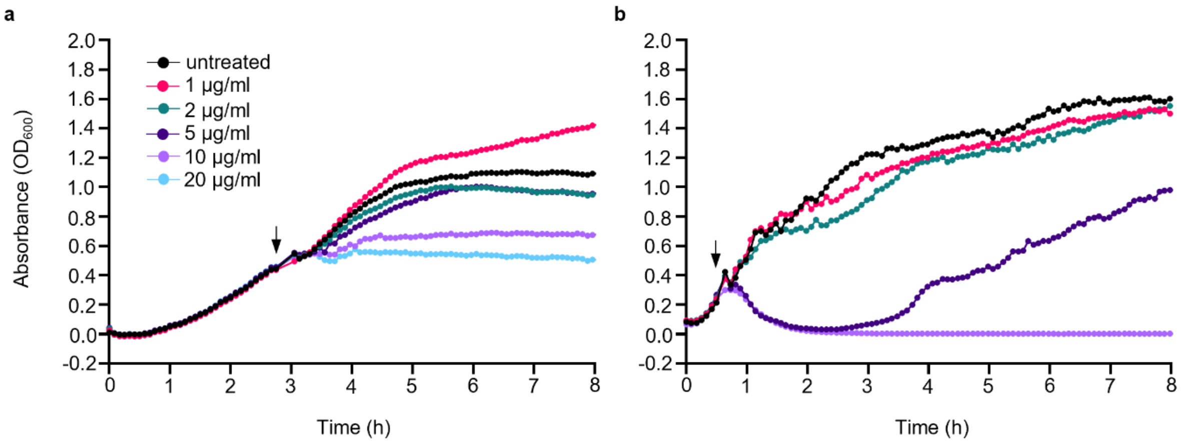


**Figure S1: Effects of daptomycin on cell growth in (a) *S. aureus* and (b) *B. subtilis*.** Growth curves of cells in LB medium supplemented with 1.25 mM $CaCl_2$ and exposed to different daptomycin concentrations at $OD_{600}$ of 0.3-0.4 (arrow). Strains used: *S. aureus* SH1000 and *B. subtilis* 168 (wild type).

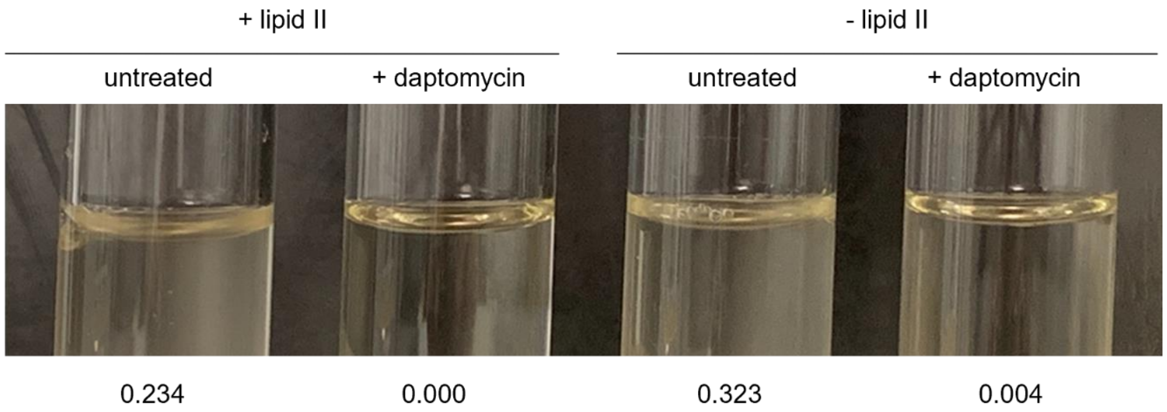


**Figure S2: Inhibition of L-form growth by daptomycin.** *B. subtilis* L-forms able and unable to produce lipid II were untreated or treated with 10 µg/ml daptomycin and incubated at 30 °C for 72 h, until visible growth was observed. Figures at the bottom of the panel represent $OD_{600}$ values. Strains used: *B. subtilis* AK092 (L-forms able to produce lipid II), *B. subtilis* AK0197 (L-forms unable to produce lipid II).

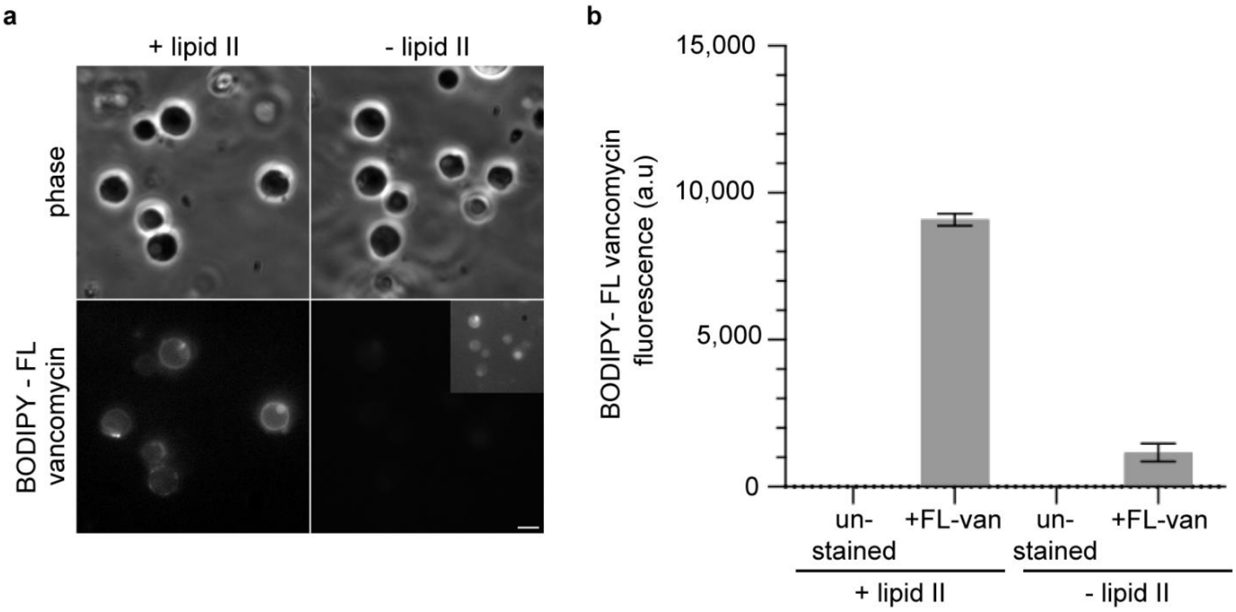


**Figure S3: *B. subtilis* AK092 L-forms still synthesise lipid II, whilst AK0197 L-forms do not.** (a) Lipid II is not detectable in the *ΔuppS* L-form mutant (- lipid II) when stained with fluorescent vancomycin (BODIPY-FL vancomycin). Phase contrast (top panel) and BODIPY-FL vancomycin (bottom panel). The large images preserve the fluorescence intensity differences between the individual conditions. The small insert images serve to visualise the presence of cells in otherwise dark image fields. Scale bar, 3µm. (b) Quantification of overall BODIPY-FL vancomycin fluorescence in a plate reader with error bars indicating the standard deviation of at least three technical replicates from one experimental set. Strains used: *B. subtilis* AK092 (L-forms able to produce lipid II), *B. subtilis* AK0197 (L-forms unable to produce lipid II).

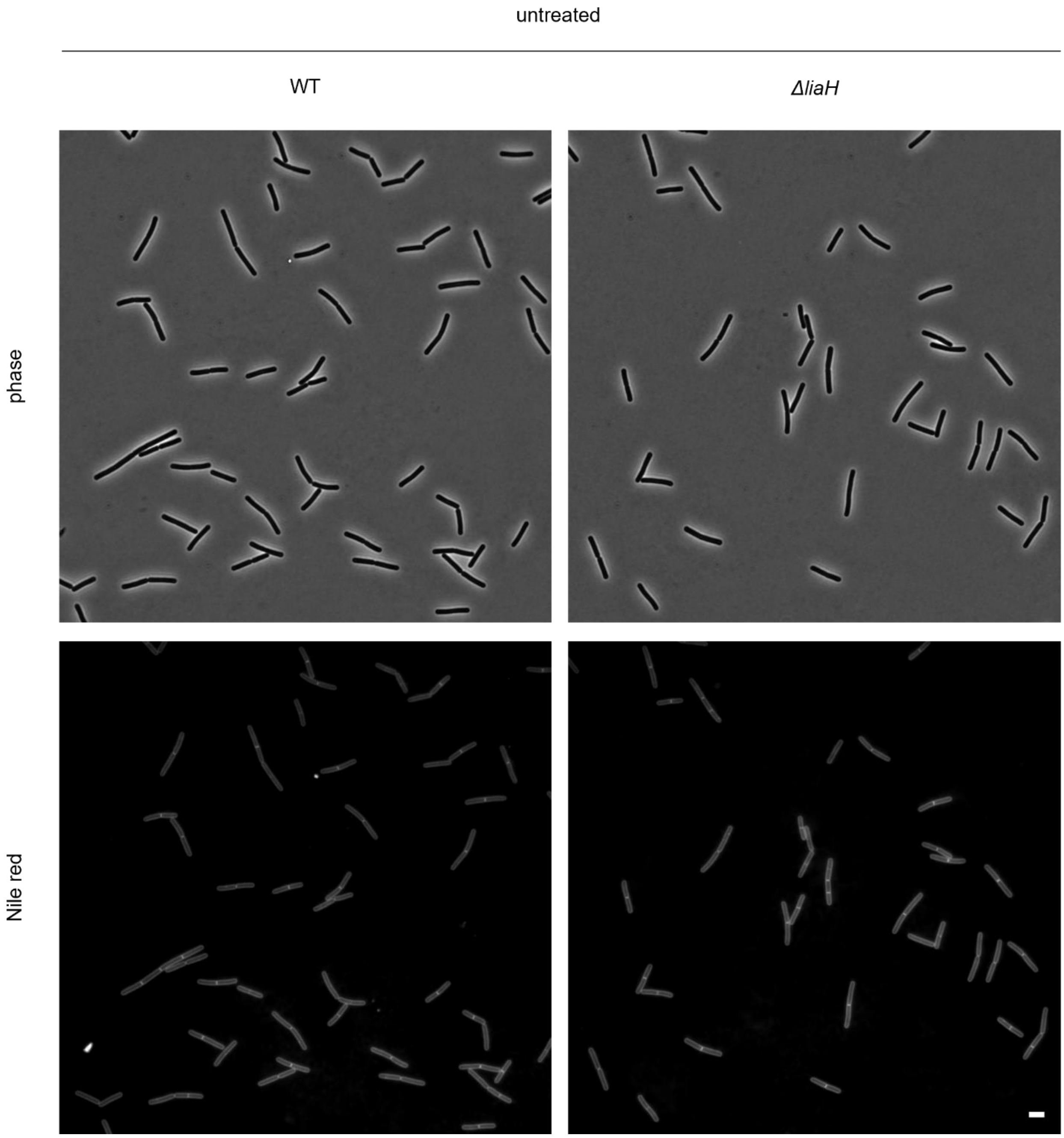


**Figure S4: Overview images of Nile red-stained untreated *B. subtilis*.** Phase contrast and fluorescence microscopy of *B. subtilis* wild type (WT) and *ΔliaH* cells stained with 1 µg/ml Nile red. Scale bar, 3µm. Strains used: *B. subtilis* 168 (wild type) and CD11 (*ΔliaH*).

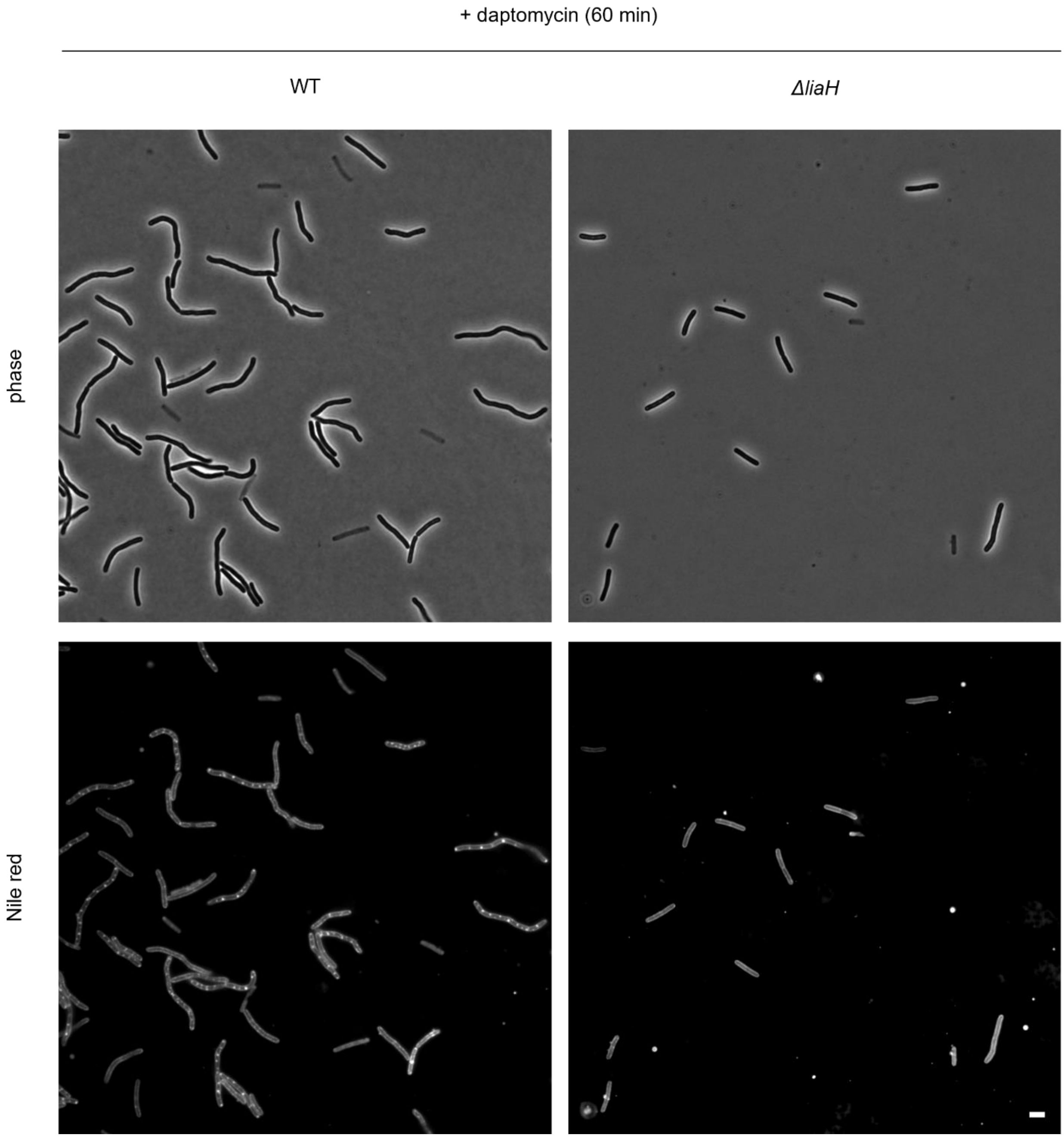


**Figure S5: Overview images of Nile red-stained daptomycin-lipid clusters.** Phase contrast and fluorescence microscopy of *B. subtilis* wild type (WT) and *ΔliaH* cells stained with 1 µg/ml Nile red and treated with 4 µg/ml daptomycin for 60 min. Scale bar, 3µm. Strains used: *B. subtilis* 168 (wild type) and CD11 (*ΔliaH*).
